# Prevalence of *Pseudomonas aeruginosa* in Australian wild birds, native wildlife, livestock and domestic animals

**DOI:** 10.1101/2025.06.23.661201

**Authors:** Kellie R. Strickland, Martina Jelocnik, Erin P. Price, Derek S. Sarovich

## Abstract

The ESKAPE pathogen, *Pseudomonas aeruginosa*, poses a serious threat to medical, veterinary, and agricultural practices globally. Understanding *P. aeruginosa* prevalence in wild bird populations, livestock, and domestic animals is vital for evaluating potential infection reservoirs. In this study, we screened 1,669 DNA samples obtained between 2010 and 2023 from healthy and diseased wild birds (*n*= 1,101), domestic animals (*n*= 269), livestock (*n*= 133), kangaroos (*n*= 39,) and koalas (*n*= 127) from Southeast Queensland, Australia, for both *P. aeruginosa* and overall bacterial load using an *ecfX*-16S rRNA duplex real-time PCR assay. *P. aeruginosa-*positive samples were also screened for the two most common fluoroquinolone resistance genotypes, GyrA Thr83Ile and GyrA Asp87Asn.

Overall, only 1.8% samples were *P. aeruginosa* positive, a lower rate than reported in international studies. Livestock samples showed the highest *P. aeruginosa* prevalence (4.5%, *n*= 6), primarily horses (7.4%, *n*= 5), with wild birds (1.5%, *n*= 17), koalas (1.6% *n*= 2), and domestic animals (1.9%, *n*= 5) having the next highest rates. In contrast, no *P. aeruginosa* positive samples were identified in cattle (*n*= 45) or kangaroos (*n*= 39). Nearly all positive wild bird samples originated from eye swabs (94%, *n*=16). No additional correlation between swab site, health status, or admission cause was identified. The GyrA Thr83Ile variant was seen in 2/30 (6.6%) *P. aeruginosa*-positive samples, both of horse origin. Our findings provide important insight into the epidemiology of *P. aeruginosa* in Australian wildlife and domestic animal populations. Further prevalence studies, particularly covering a broader geographical region, are warranted to better elucidate nationwide *P. aeruginosa* carriage, infection, and fluoroquinolone resistance rates.

## Introduction

The environmentally ubiquitous and moisture-loving bacterium, *Pseudomonas aeruginosa*, poses a significant and increasing burden across medical and veterinary industries (1,2). Its high intrinsic resistance towards many antibiotics, extreme genetic adaptability, and ability to form tenacious biofilms make this opportunistic pathogen particularly problematic for those with chronic respiratory conditions, such as cystic fibrosis, often resulting in irreversible lung decline and early death in susceptible cohorts (3,4). *P. aeruginosa* can also cause a plethora of diseases in companion animals, livestock and wildlife, where it typically presents as otitis media, ulcerative keratitis, and urinary tract infections in cats and dogs, mastitis in dairy cows, endometritis, genital, respiratory tract, and eye infections in horses, and pneumonia, septicaemia, keratitis, and embryonic death in birds and poultry (5). Due to its significant morbidity and mortality impacts, the potential for transmission between different hosts, and its global ubiquity, tackling *P. aeruginosa* requires a strong ‘One Health’ approach.

Anthropogenic activities have led to environments such as farmlands, sewage facilities, and waterways becoming contaminated with antimicrobial-resistant (AMR) bacteria, which subsequently can infect the livestock and wildlife that inhabit them (6–8). For instance, wild birds can act as a potential reservoir and vector for AMR bacteria, resulting in further environmental dissemination that increases human infection risk (6–9). This phenomenon has been documented in Sweden (2017) and Australia (2016), whereby up to 40% of wild gulls harboured either extended-spectrum β-lactamase (ESBLs), or carbapenemases producing *Escherichia coli,* respectively. Interestingly, these isolates demonstrated highly similar genetic characteristics with local human clinical isolates, indicating probable zoonotic transmission (10,11). Multidrug-resistant *Pseudomonas* species have been isolated from wild bird faeces near the River Cam, England, with an alarming 85% expressing resistance towards the first-line antibiotic, ciprofloxacin (12). In Lebanon, VIM-2 carbapenemase-producing *P. aeruginosa* have been isolated from pigs, poultry, and cattle (13). Alarmingly, *P. aeruginosa* carrying mobile colistin resistance genes *mcr-1* & *mcr-2* have been discovered in resident and migratory wild birds in Egypt, indicating potential widespread dissemination of this AMR gene, which confers resistance towards this critical last-line antibiotic (14). *P. aeruginosa* carrying *mcr* has not yet been documented in Australia; however, in 2019, *mcr-1-* encoding *E. coli* was isolated from a silver gull, the first time this AMR gene family has been identified in Australian animals (15). Together, these studies highlight the potential risk of unmitigated and widespread AMR *P. aeruginosa* dissemination from the environment to humans and animals.

International studies of wild bird populations, livestock, and domestic animals have found that *P. aeruginosa* prevalence can vary widely, although there are some parallels based on the animal cohort, sample location, and animal health status (5,16). For instance, *P. aeruginosa* prevalence is relatively consistent (4.6 – 11.7%) in the horse urogenital tract (17–20), but exhibits more variability (11.8 – 64.3%) in horse ocular infections (20,21). Less is known regarding companion animals, although *P. aeruginosa* is considered to be more common in dogs than cats, whereby *P. aeruginosa* prevalence ranges from 4.7–18.7% in dogs (22–24), and 1.2–5.2% in cats (5,22,25). Livestock incidence is variable but tends to be highest in poultry, with rates between 8.2 and 52% being reported (26,27). Recent screening of faecal and cloacal samples from wild birds found that *P. aeruginosa* prevalence ranged from 7.8– 11.1% in Spain and Italy (28–30) to as high as 20% in Egypt and England (12,14).

There is a comparative dearth of research on *P. aeruginosa* prevalence in Australian animals. Most of Australia’s wildlife is found nowhere else in the world, with 45% of bird, 93% of reptile, and 87% mammal species classed as endemic (9,31,32). To address this knowledge gap, we screened 1,669 retrospective DNA samples from a diverse range of wild and domestic animals in Southeast Queensland, Australia, using a duplex real-time PCR assay targeting *P. aeruginosa* and overall bacterial load.

## Methods

### Ethics approval

All study samples were deemed as exempt from animal ethics requirements, as determined by the University of the Sunshine Coast Animal Ethics committee (waivers ANE1939, ANE1940, and ANE2057).

### DNA samples

This study examined genomic DNA of 1,669 retrospective samples collected between 2010 and 2023 from healthy and diseased animals residing in Southeast Queensland, Australia (33,34). Approximately two-thirds of these samples (*n*=1,101) originated from 565 euthanised wild birds sampled opportunistically during admission at the Australia Zoo Wildlife Hospital (AZWH, Beerwah, Queensland) between May 2019 and December 2021 (565 eye, 534 cloacal, and two “other” swabs) encompassing 112 species and 44 families, predominantly parrots (Psittaculidae: 17.2%), pigeons and doves (Columbidae: 14.3%), cockatoos (Cacatuidae: 11.7%), kingfishers (Alcedinidae: 8.7%), honeyeaters (Meliphagidae: 6.9%), and birds of prey (6.8%) (33,34) (**Table S1**). Most wild birds were native to Australia (96.9%). These birds were originally admitted due to trauma (53.2%), clinical disease (23.2%), animal attacks (9.7%), or other miscellaneous causes (2.1%); however, some were not specified (11.8%).

This study included retrospective DNA samples from domestic animals (*n*=269 samples; 138 cat, 126 dog, 3 pet guinea pig, 1 pet bird, 1 pet rabbit), livestock (*n*=133 samples; 68 horse, 45 cattle, 14 goat, 3 alpaca, 3 poultry), koala (*n*=127), and kangaroo (*n*=39) samples that were opportunistically acquired at AZWH, local veterinarian centres, or within their native habitat (e.g. faecal samples) (**Table S2**) (35,36). Some samples were acquired from the same animal (e.g. all 14 goat samples were from one necropsied animal), although most samples were from individual animals. DNA samples were sourced from a variety of anatomical sites, including eye (*n*= 623), cloaca (*n*=537), faeces (*n*=314), respiratory tract (*n*=62), female reproductive tract/organs (*n*=59), urogenital tract (*n*=45), and rectum (*n*=20), along with several other less commonly sourced sites such as joint fluid (*n*=7), fur (*n*=2), penis (*n*=2), intestine (*n*=1), liver (*n*=1), brain (*n*=1), prepuce (*n*=1), stifle (*n*=1), spleen (*n*=1), and others (**Table S2**).

### Quantitative real-time PCR

All DNA samples were screened in duplicate for *P. aeruginosa* presence and overall bacterial load using an *ecfX* (*PA1300*)-16S rRNA duplex real-time PCR assay targeting *P. aeruginosa* (37) and all bacteria (38), respectively (**Table 1**). Reactions were carried out in opaque, white, hardshell 96-well PCR plates (Bio-Rad, Gladesville, NSW, Australia) using Microseal C PCR plate sealing film (Bio-Rad) and the CFX96 Touch real-time PCR instrument (Bio-Rad). 1 µL DNA was added to a 5 µL total reaction volume, using qPCRBIO Probe mix (PCR Biosystems, London, UK), unlabelled primers (IDT, Coralville, IA, USA), and Black Hole Quencher probes (LGC Biosearch Technologies, Petaluma, CA, USA) (**Table 1**). Thermocycling conditions comprised an initial denaturation at 95°C for 2 min, followed by 45 cycles of 95°C for 10 sec and 60°C for 15 sec. For each plate, at least one positive control (*P. aeruginosa* strain SCHI0005.S.8 (39); NCBI ID: SRR15793216) and one reagent-only no-template control (NTC) were included to ensure mastermix integrity. All data were analysed with CFX Maestro software using the ‘Regression’ threshold setting to permit inter-run comparisons.

**Table 1.**
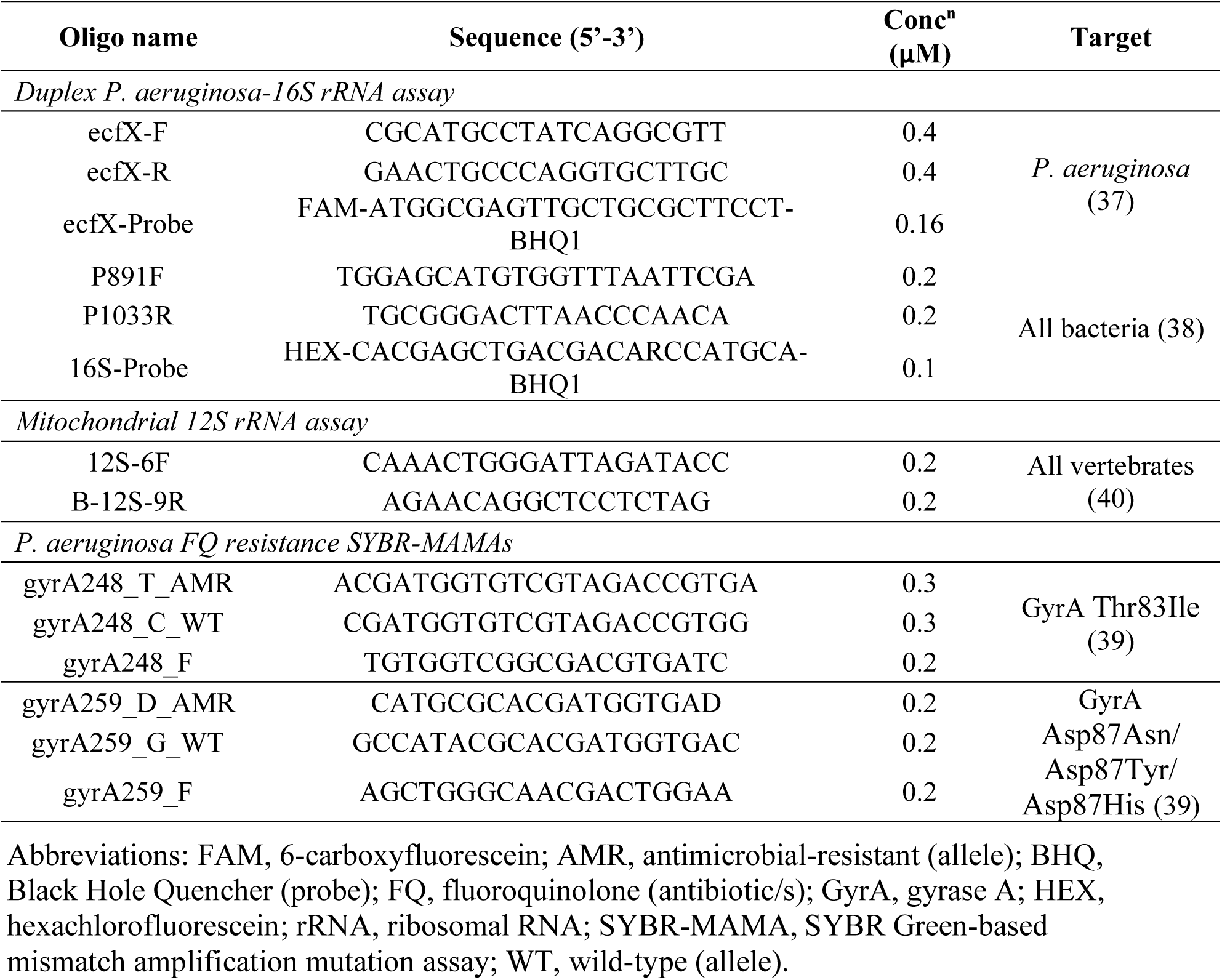
Real-time PCR primers, probes, and assays used in this study.

For each sample, pan-bacterial cycles-to-threshold (*C*_T_) values were first assessed to ensure DNA integrity and to confirm bacterial DNA presence. The cut-off for successful pan-bacterial amplification above background was determined by calculating the average NTC *C*_T_ value across 10 separate runs, minus one standard deviation. Using this cut-off metric, samples with a repeatable 16S *C*_T_ < 25.9 (as determined by a minimum of two replicates) were considered to possess sufficient bacterial DNA for *P. aeruginosa* (*ecfX*) assessment. In 116 samples that did not meet this pan-bacterial threshold, or that failed to generate robust amplification curves across replicates, the DNA was quantified and assessed using a NanoDrop spectrophotometer (Thermo Scientific, Scoresby, VIC, Australia) and Qubit 3 fluorometer (Thermo Scientific), then diluted to 2 ng/μL (if >20 ng/μL) and repeated.

Overall, a total of 1700 samples were screened, but 31 samples failed the pan-bacterial threshold on repeat testing and were excluded from the study due to poor DNA quality, resulting in 1669 valid DNA samples.

Once DNA quality was confirmed, we next assessed *P. aeruginosa* positivity in the 1,669 samples. Similar to the pan-bacterial assay, the upper *P. aeruginosa ecfX-*positive *C*_T_ cut off was determined by calculating one standard deviation range of the NTC amplification determined from 10 separate runs. Using this cut-off, samples with a repeatable *ecfX C*_T_ < 35.7 were considered positive.

To better understand pathogen-to-host load in *P. aeruginosa (ecfX)* positive samples and sample suitability for metagenomic sequencing, a mitochondrial 12S rRNA assay that amplifies all vertebrates (40) was next tested (**Table 1**). These PCRs, which consisted of 1X SsoAdvanced SYBR Green Mix (Bio-Rad) and 1µL template to a 5uL total reaction volume, were undertaken using previously described thermocycling parameters (41), except that initial denaturation was for 2 min only, and there was no final extension. All samples amplified successfully and before the average NTC *C*_T_ of 36.5.

### AMR genotyping

All *P. aeruginosa* positive samples were screened for two fluoroquinolone resistance genotypes (GyrA Thr83Ile and GyrA Asp87Asn, Asp87Tyr, and Asp87His) using SYBR Green-based mismatch amplification mutation assays (SYBR-MAMAs) (**Table 1**) (39). These mutations are responsible for the majority of fluoroquinolone AMR seen in clinical *P. aeruginosa* isolates (39,42,43). *P. aeruginosa* SCHI0002.S.8 (NCBI ID: SRR15793214) was used as a wild-type (WT) strain for GyrA

Thr83Ile while SCHI0004.S.5 (NCBI ID: SRR15793159) was the WT strain for GyrA Asp87Asn. SCHI0033.S.15 (NCBI ID: SRR15793165) and SCHI0005.S.9 (NCBI ID: SRR15793213) were used as AMR strains for GyrA Thr83Ile and GyrA Asp87Asn, respectively (39). Thermocycling conditions were adjusted to 40 cycles, and used an extended two-step program (95°C for 10 sec and 60°C for 15 sec), to allow adequate annealing and extension time for polymicrobial DNA samples.

## Results

### *P. aeruginosa* prevalence in Australian veterinary samples

Overall *P. aeruginosa* prevalence in Australian animal samples was low, being < 8% among all animal categories with adequate sample size (i.e. *n=* >20), and 1.8% across all 1,669 tested samples (**Table 2**). Livestock showed the highest *P. aeruginosa* prevalence (4.5%, *n*=6), driven primarily by horses (7.4%, *n*=5). Next most prevalent were wild birds (1.5%, *n*=17), koalas (1.6%, *n*=2), and domestic animals (1.9%, *n*=5), which was primarily composed of cats (2.2%, *n*=3). No positive samples were identified in samples from cattle (*n*=45), kangaroo (*n*=39), goat (*n*=14), guinea pig (*n*=3), poultry (*n*=3), or rabbit (*n*=1) (**Table 2**).

**Table 2.** Overview of all veterinary DNA samples tested in this study, and their *Pseudomonas aeruginosa* positivity rates.

|  | No. samples (%<br>total) | No. *P. aeruginosa*<br>positive (%) |
| --- | --- | --- |
| **Wild birds** | 1101 (66.0) | 17 (1.5) |
| **Livestock** |  |  |
| Horse | 68 (4.1) | 5 (7.4) |
| Cattle | 45 (2.7) | 0 (0) |
| Goat | 14 (0.8) | 0 (0) |
| Alpaca | 3 (0.2) | 1 (33) |
| Poultry | 3 (0.2) | 0 (0) |
| **Native marsupials** |  |  |
| Koala | 127 (7.6) | 2 (1.6) |
| Kangaroo | 39 (2.3) | 0 (0) |
| **Domestic animals** |  |  |
| Cat | 138 (8.3) | 3 (2.2) |
| Dog | 126 (7.5) | 1 (0.8) |
| Pet bird | 1 (0.1) | 1 (100) |
| Guinea Pig | 1 (0.1) | 0 (0) |
| Rabbit | 1 (0.1) | 0 (0) |
| **TOTAL** | 1669 | 30 (1.8) |

Nearly all *P. aeruginosa-*positive wild bird samples originated from eye swabs (94.2%, *n*=16/17), with only one positive cloaca sample (5.8%, n=1/17). All horse penile samples (*n*=2) and tracheal washes (*n*=2) had *P. aeruginosa*; however, due to very low sample size, the significance of these infection sites could not be ascertained. No other additional correlations between swab site, health status, or admission cause were identified (**Table 3**).

**Table 3.** Summary of all *Pseudomonas aeruginosa* PCR-positive samples identified in Australian animals.

| Sample ID | Animal | Common species name | Admission cause | Site |
| --- | --- | --- | --- | --- |
| 90947E | Wild bird | Yellow-tailed Black-Cockatoo | Clinical disease (emaciated) | Eye |
| 93645E | Wild bird | Little Corella | Unknown | Eye |
| 101198E | Wild bird | Sulphur-crested Cockatoo | Clinical disease (BFD) | Eye |
| 93618E | Wild bird | Rainbow Lorikeet | Clinical disease (unsp.) | Eye |
| 94279E | Wild bird | Rainbow Lorikeet | Clinical disease (poxvirus) | Eye |
| 94403E | Wild bird | Crested Pigeon | Clinical disease (emaciated) | Eye |
| 87321E | Wild bird | Australian Pelican | Clinical disease (unsp.) | Eye |
| 87187E | Wild bird | Australian Magpie | Clinical disease (ocular) | Eye |
| 87154E | Wild bird | Spangled Drongo | Animal attack (unsp.) | Eye |
| 87930E | Wild bird | Australian Brush-turkey | Animal attack (unsp.) | Eye |
| 86932C | Wild bird | Noisy Miner | Clinical disease (ocular) | Cloaca |
| 88732E | Wild bird | Australasian Figbird | Trauma (unsp.) | Eye |
| 90083E | Wild bird | Masked Lapwing | Trauma (HBC) | Eye |
| 25102019E | Wild bird | Noisy Miner | Unknown | Eye |
| 90971E | Wild bird | Pied Butcherbird | Trauma (unsp.) | Eye |
| 310120202E | Wild bird | Black-faced Cuckoo-shrike | Clinical disease (poxvirus) | Eye |
| 101493E | Wild bird | Eastern Koel | Trauma (unsp.) | Eye |
| av conj 80 | Bird | Domestic (unsp.) | Unknown/opportunistic | Eye |
| 50F | Cat | Domestic Cat | Unknown/opportunistic | Faeces |
| ALP rec | Alpaca | Huacaya Alpaca | Unknown/opportunistic | Rectum |
| sphnx cat oral | Cat | Domestic Cat | Unknown/opportunistic | Oral |
| sphnx conj | Cat | Domestic Cat | Unknown/opportunistic | Eye |
| KP5 * | Horse | Domestic Horse | Unknown | Faeces |
| TRCIC 082H | Horse | Domestic Horse | Unknown | Tracheal wash |
| Trach 882 * | Horse | Domestic Horse | Unknown | Tracheal wash |
| S50 penile | Horse | Domestic Horse | Unknown | Penile |
| S79 penile | Horse | Domestic Horse | Unknown | Penile |
| 56 | Dog | Domestic Dog | Unknown | Faeces |
| 1 O | Koala | Wild Koala | Unknown | Eye |
| 42 Branson | Koala | Wild Koala | Nasal Discharge | Nasal |

\* Potential fluoroquinolone-resistant infection based on *gyrA248* genotyping (see Figure 2).
Abbreviations: BFD: beak and feather disease; HBC: hit by car; unsp.: Unspecified.

### Mitochondrial 12S pan-vertebrate real-time PCR testing in *P. aeruginosa* samples

All 30 *P. aeruginosa-*positive samples were found to have high burden of host DNA present within samples. Due to the retrospective nature of these samples, host DNA depletion was not possible, and therefore none of our *P. aeruginosa-*positive samples were able to undergo metagenomic sequencing.

### Fluoroquinolone AMR in *P. aeruginosa-*positive samples

All 30 *P. aeruginosa-*positive samples harboured the *gyrA259* wild-type (WT) genotype (**Figure 1**). In contrast, 2/30 (6.6%) samples, both from horses (KP5 and Trach 882; **Table 3**), possessed evidence of a mixed *gyrA248* phenotype (i.e. both WT and AMR variants present) based on much smaller Δ*C*_T_ values compared with control WT and AMR strains (**Figure 2**). This observation suggests polymicrobial infections containing both fluoroquinolone-sensitive and -resistant *P. aeruginosa* in these two horses.

**Figure 1.**
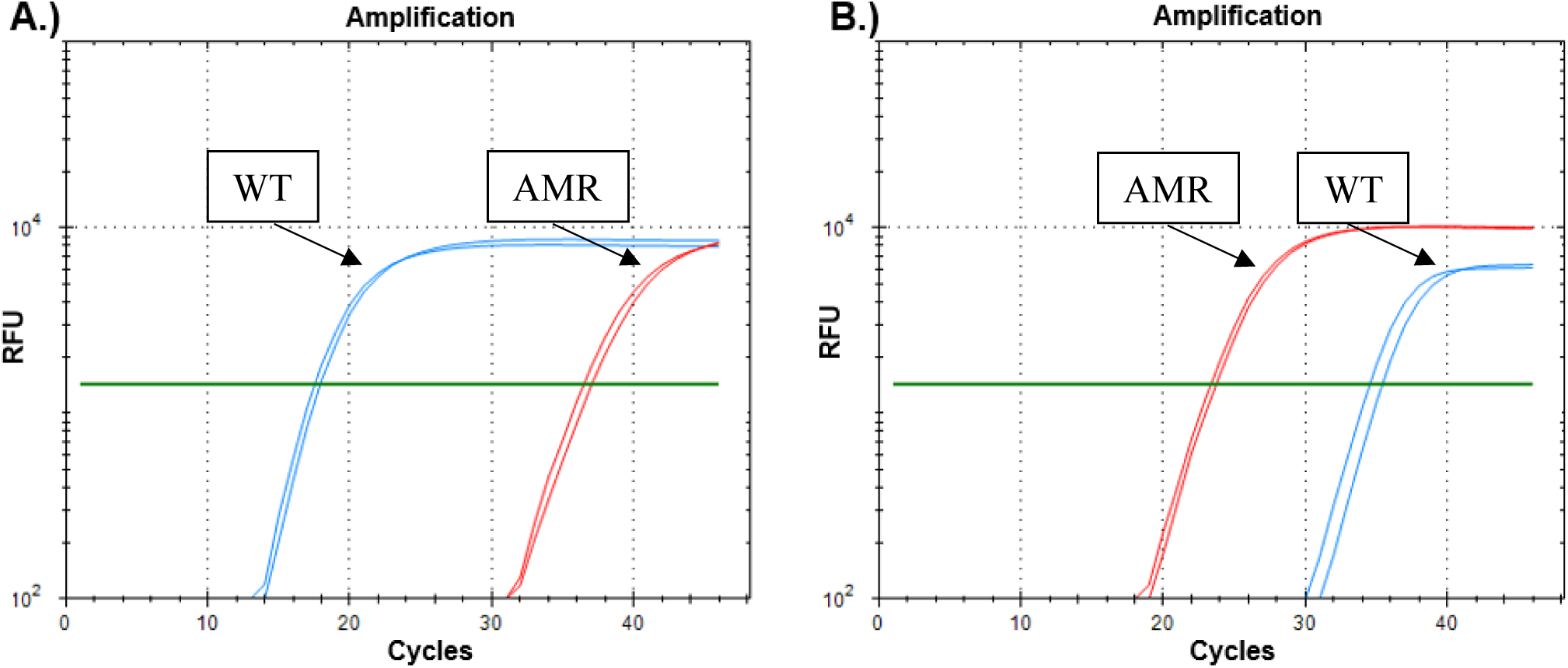
GyrA Asp87Asn/Tyr/ His SYBR Green-based mismatch amplification mutation assay targeting the Pseudomonas aeruginosa fluoroquinolone resistance-conferring tetra-allelic single-nucleotide polymorphism at gyrA259. (39) (A) Wild type (WT) gyrA259 control strain SCHI0004.S.5 and (B) antimicrobial-resistant (AMR) control strain SCHI0005.S.9. Blue line=WT allele, red line=AMR allele. All P. aeruginosa veterinary samples unambiguously exhibited the WT gyrA259 genotype. RFU, relative fluorescence units.

**Figure 2.**
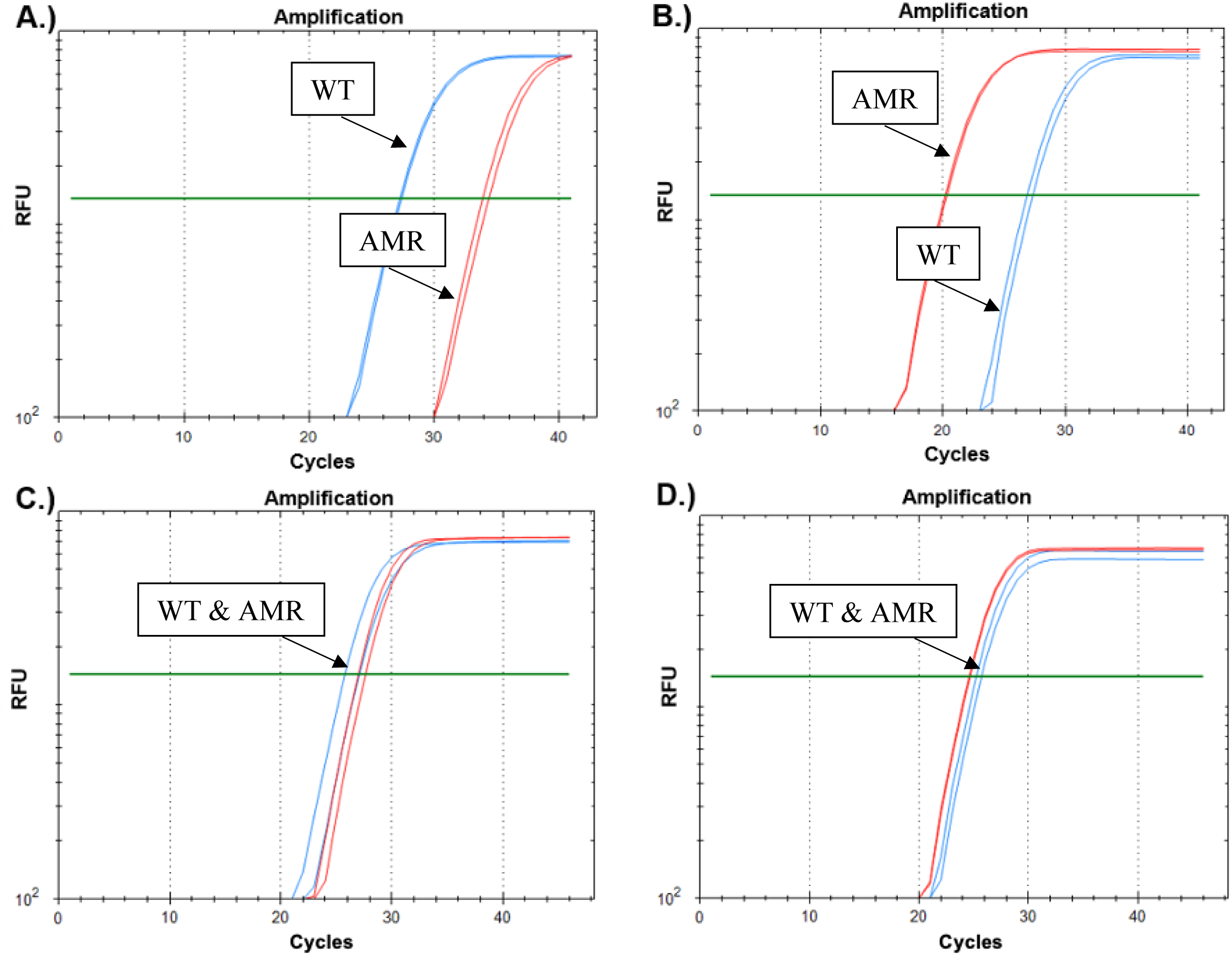
GyrA Thr83Ile SYBR Green-based mismatch amplification mutation assay targeting the *Pseudomonas aeruginosa* fluoroquinolone resistance-conferring single-nucleotide polymorphism at *gyrA*248. **(A)** Wild type (WT) *gyrA*248 control strain SCHI0002.S.8 and **(B)** antimicrobial-resistant (AMR) control strain SCHI0033.S.15. Blue line=WT, red line=AMR. All *P. aeruginosa* veterinary samples unambiguously exhibited the WT *gyrA248* genotype, except for horse samples 882trach **(C)** and KP5 **(D)**, which possessed a mixed *gyrA248* phenotype (i.e. both WT and AMR variants were present at approximately equal abundance). RFU, relative fluorescence units.

## Discussion

This study determined the prevalence of the ESKAPE pathogen, *P. aeruginosa,* across a large spectrum of both healthy and diseased animals residing in subtropical Southeast Queensland, Australia. Overall, only 1.5% of samples from our large and diverse sample set tested positive for *P. aeruginosa*.

To the best of our knowledge, our study is the first to investigate *P. aeruginosa* prevalence in Australian wild birds. Prior studies have focused on *E. coli* or *Salmonella* spp. prevalence in wild Australian birds, predominantly gulls, as these synanthropic birds act as ecological sponges and AMR gene collectors (11,15,44–46). Specifically, 504 gulls from an Australian nesting site displayed a high positivity rate for salmonellae (30%) and *E. coli* (40%), with an alarming 50.8% of these *E. coli* isolates carrying *bla*_IMP_ carbapenemase genes (11). In contrast, all 22 gulls we tested were negative for *P. aeruginosa*, suggesting that gulls may not play an important role in *P. aeruginosa* carriage, AMR, and transmission in our region, although broader testing efforts would be needed to confirm this hypothesis. Overall, we found *P. aeruginosa* prevalence among wild birds was 1.5% (17/1,101), which correlates well with the 2% prevalence rate of *Pseudomonas* spp. from a 1988 American study by Brittingham *et al*. of cloacal swabs from 364 opportunistically caught wild birds (47).

However, our rate is substantially lower than the rates reported in wild birds from Italy (11.1%; 18/163) (27), in resident (10%; 8/80) and migratory (18.3%; 11/60) birds from Egypt (14), and in diseased wild raptors in Spain (7%; 32/457) (14,28,30). Furthermore, our positive *P. aeruginosa* samples were essentially all ocular (94%) in origin, with only one positive cloacal sample found. This is in stark contrast to Russo *et al*., who investigated only cloacal samples in the Italian wild bird cohort, yet observed an 11.1% *P. aeruginosa* positivity rate (30). Of note, most *P. aeruginosa* isolated from wild Spanish raptors by Vidal *et al*. were from birds with systemic infection, respiratory illness, or oral lesions, with none isolated from birds with conjunctivitis or diarrhoea (28); therefore, the rates of *P. aeruginosa* infection in these bird populations may be even higher. Our lower *P. aeruginosa* prevalence rate in wild birds is likely due to our inclusion of healthy birds and swabbing across multiple anatomical sites. Based on our broader sampling efforts, our results may better reflect the pathogen’s occurrence across wild bird populations.

Although relatively uncommon, *P. aeruginosa* is a known opportunistic pathogen of koalas, having been documented as early as 1978 as the causative agent behind a fatal epidemic in koala pouch joeys (48). However, aside from a 1986 case report of fatal koala bronchopneumonia (49), and a small-scale examination of the koala pouch microflora in 1992 (50), almost no research has been undertaken to determine *P. aeruginosa* prevalence in koalas. Among 127 koala samples, we found a *P. aeruginosa* prevalence rate of 1.6% (*n*=2), whereby one eye and one nasal swab were positive. Our results support the findings from the National Koala Disease Risk Analysis Report (2023) that acknowledges *P. aeruginosa* disease risk potential in Australian koalas (51), albeit not at significant levels.

*P. aeruginosa* is thought to have a lesser burden in other native Australian mammals such as kangaroos, with minimal disease burden reported in adults, although *P. aeruginosa* can cause peritonitis or pneumonia in joeys (52,53). We found no positive *P. aeruginosa* faecal samples from 39 adult kangaroos, supporting this view. In stark contrast, a smaller Australian study found 42.9% (6/14) of kangaroo faecal samples contained *P. aeruginosa* based on a *regA* (*PA0707*; also called *toxR*)-based SYBR Green-based real-time PCR assay, the greatest prevalence seen in any of the various animals tested (54). The basis for this difference in *P. aeruginosa* positivity rates in Southeast Qld kangaroo populations is unknown, but may be due to non-specific *regA*/*toxR* amplification in non-*aeruginosa* pseudomonads such as *P. paraeruginosa*.

*P. aeruginosa* has a greater disease burden in companion animals than in wild animals, potentially posing a zoonotic threat for immunocompromised people with pets (5,55,56). One Italian study found that 8% (24/300) clinically unwell dogs carried *P. aeruginosa*, 25% of which had ear infections (23). A South Korean study reported an 18.7% (84/448) positivity rate, with no significance difference between healthy (17.7%) and unhealthy (16.7%) dogs (24). A similarly high *P. aeruginosa* rate was also seen in a small Australian study (14.3%), although only 14 dog faecal samples were tested (54). The lowest *P. aeruginosa* prevalence rates reported in companion dogs to date was from a large Thai study, which found *P. aeruginosa* in 4.7% of unwell dogs from a variety of sample types (22). In contrast, our findings were significantly lower than these prior canine studies, with a *P. aeruginosa* prevalence of just 0.8% in Australian dogs (1/126). The discrepancy in prevalence rates is likely due to the opportunistic nature of our canine faecal samples, whereas most of the aforementioned studies targeted dogs with signs of clinical disease typical of *P. aeruginosa*, such as otitis externa and eye infections. Unexpectedly, we found a *P. aeruginosa* prevalence rate of 2.2% (3/138) in domestic cat samples, despite this pathogen typically being more prevalent in dogs (5). Our rate is higher than that reported in the Thai study (1.2%) but lower than a Spanish study (5.2%), both of which investigated only unwell cats and various anatomical sites (22,25). Taken together, there appears to be minimal, but not zero, risk of zoonotic *P. aeruginosa* transmission between companion animals and their owners in Australia. Further studies are needed to resolve prevalence rates in domestic cats in our region.

Globally, *P. aeruginosa* is an important venereal disease and major cause of endometritis in horses, resulting in infertility or early gestational failure (20,57). Accordingly, pre-breeding screening for *P. aeruginosa* is strongly advised by the International Codes of Practice declared by the UK’s Horserace Betting Levy Board (HBLB), which tests >20,000 samples per annum in the United Kingdom alone (58). As a part of this screening, between October 2019 and December 2024 nearly 30,000 samples underwent *P. aeruginosa* PCR screening, from which just 159 (0.5%) were positive (58,59). A large-scale review of studies performed between 1996 and 2021 in various countries such as the UK, United States, Europe, and the United Arab Emirates found that *P. aeruginosa* prevalence in horse uterine infections was 0 – 11.7% (20). Our study falls within this range, with 7.4% (5/68) *P. aeruginosa*-positive horse samples from faeces (1/35), respiratory tract (2/25), and penile swabs (2/2). Our findings are marginally elevated compared to Moore and colleagues, who found *P. aeruginosa* in 5.4% in 280 horses in Ireland (18), and Kidd and colleagues, who found 4.6% *P. aeruginosa* in 2,040 mares and 18 stallions across 66 thoroughbred horse stud farms in Southeast Queensland (19), the same geographic region as our study. Among the Kidd *et al*. positive samples were mare genital swabs (4.6%), 45% of which were clonally related strains of a single dominant genotype (19), whereas none of the female genital swabs tested in our study (*n*=21) possessed *P. aeruginosa*. Further, none of the 18 stallions studied by Kidd *et al*. were *P. aeruginosa* positive; in comparison, our two horse penis swabs were both positive. These discordant results may be due to different horse populations (thoroughbreds vs. domestic pets), sampling times (2005-2009 vs. 2010-2023) or may simply reflect our study’s lower sample numbers. Overall, *P. aeruginosa* prevalence detected in horses in our study correlates well with other studies conducted both nationally and internationally.

Fluoroquinolones are categorised by the World Health Organisation as highest priority, critically important antimicrobials, and as such their use is restricted in all animals, including horses (60). The British Equine Veterinary Association and Equine Veterinarians Australia both recommend limiting fluoroquinolone use only for cases where no alternative antibiotic choice exists, and ideally with corresponding evidence of pathogen sensitivity on antibiotic susceptibility testing (61,62). Despite these restrictions, a large study by Dorph *et al*. of 8,492 horses at four racetracks in the United States revealed the most frequently prescribed antimicrobial was enrofloxacin (31.8%; *n*=854), and alarmingly, this fluoroquinolone was used as first-line treatment or prophylaxis in most cases (63). Kidd *et al*. found that 80% (74/93) of *P. aeruginosa* isolated from Southeast Qld thoroughbred mares between 2005- 2009 showed decreased susceptibility or complete resistance to enrofloxacin (19). In Sydney, Australia, a smaller study of *P. aeruginosa* from 38 horses with ulcerative keratitis found a lower but still substantial 50% enrofloxacin resistance rate (21). Internationally, Pottier *et al*. and Isgren *et al*. both found a lower *P. aeruginosa* fluoroquinolone AMR rate of 19% and 23% among 135 and 286 horses, respectively (17,64). Only Pottier *et al*. undertook molecular analysis of their strains to identify the prevalence of three fluoroquinolone AMR determinants: CrpP (a putative ciprofloxacin-modifying enzyme (17,65)), QnrVC1, and ParE Ala473Val. We have recently shown that neither CrpP presence nor the ParE Ala473Val variant are significantly associated with fluoroquinolone AMR phenotypes in *P. aeruginosa* (66), whereas two GyrA mutations, Thr83Ile and Asp87Asn, are found in up to 83% of Australian fluoroquinolone resistant strains (39). We identified the GyrA Thr83Ile variant in 2/30 (6.6%) *P. aeruginosa*-positive samples, albeit both as mixtures that also contained WT strains. Both Thr83Ile positive samples were from horses. To the best of our knowledge, this is the first reported instance of this clinically important fluoroquinolone AMR mutation in *P. aeruginosa* isolated from horses. However, the GyrA Thr83Ile variant has been found in *P. aeruginosa* isolates from sick dogs and cats from multiple studies (22,24,67); and recently another fluoroquinolone resistance conferring gene, *qnrVC1*, was isolated from MDR *P. aeruginosa* strains from donkeys in China (68). Taken together, inappropriate veterinary fluoroquinolone use and overuse may be driving *P. aeruginosa* AMR development towards this critically important drug class in many countries, including in Australia. Further investigations are needed to evaluate the true prevalence of fluoroquinolone AMR phenotypes and genotypes in both domestic and wild animals, and whether further restrictions on fluoroquinolone use in veterinary settings are warranted.

We acknowledge several study limitations. Due to the retrospective and opportunistic nature of the DNA samples used in this study, the sample set predominantly consisted of faecal and cloacal samples; more clinically relevant sources such as ocular, respiratory, or equine urogenital tract samples would likely have yielded higher *P. aeruginosa* positivity rates. Due to our focus on bird populations, some animal groups were insufficiently sampled, potentially skewing our *P. aeruginosa* prevalence rates. As all specimens were destroyed during the DNA extraction process, microbial culture could not be undertaken on any of the *P. aeruginosa*-positive samples, prohibiting phenotypic confirmation of fluoroquinolone AMR variants through antibiotic susceptibility testing. Finally, high host DNA burden, as determined by the mitochondrial 12S rRNA qPCR assay (40), meant that metagenomic sequencing of *P. aeruginosa*-positive specimens was infeasible. As such, we were unable to determine *P. aeruginosa* sequence types, uncover epidemiological links to environmental and clinical isolates, or comprehensively investigate the presence of other AMR determinants.

## Conclusion

This study provides important insights into *P. aeruginosa* prevalence across diverse animal populations in Southeast Qld, Australia, a geographical location renowned for its unique endemic wildlife. With an overall detection rate of 1.5%, our findings suggest lower *P. aeruginosa* prevalence in this region compared to international studies. Livestock, mainly horses, exhibited the highest positivity rates, followed by wild birds and domestic animals, with wild native marsupials having the lowest prevalence. The novel detection of fluoroquinolone resistance genotype GyrA Thr83Ile in horse-derived samples underscores the need for improved antimicrobial stewardship and ongoing surveillance of AMR in animal populations, particularly because of the zoonotic potential of this pathogen. Expanding geographic coverage and incorporating additional animal species in future studies will be essential to fully understand the distribution, infection rates and AMR patterns of *P. aeruginosa* in Australian wildlife, livestock, and domestic animals.

## Supplementary Tables

**Table S1.**
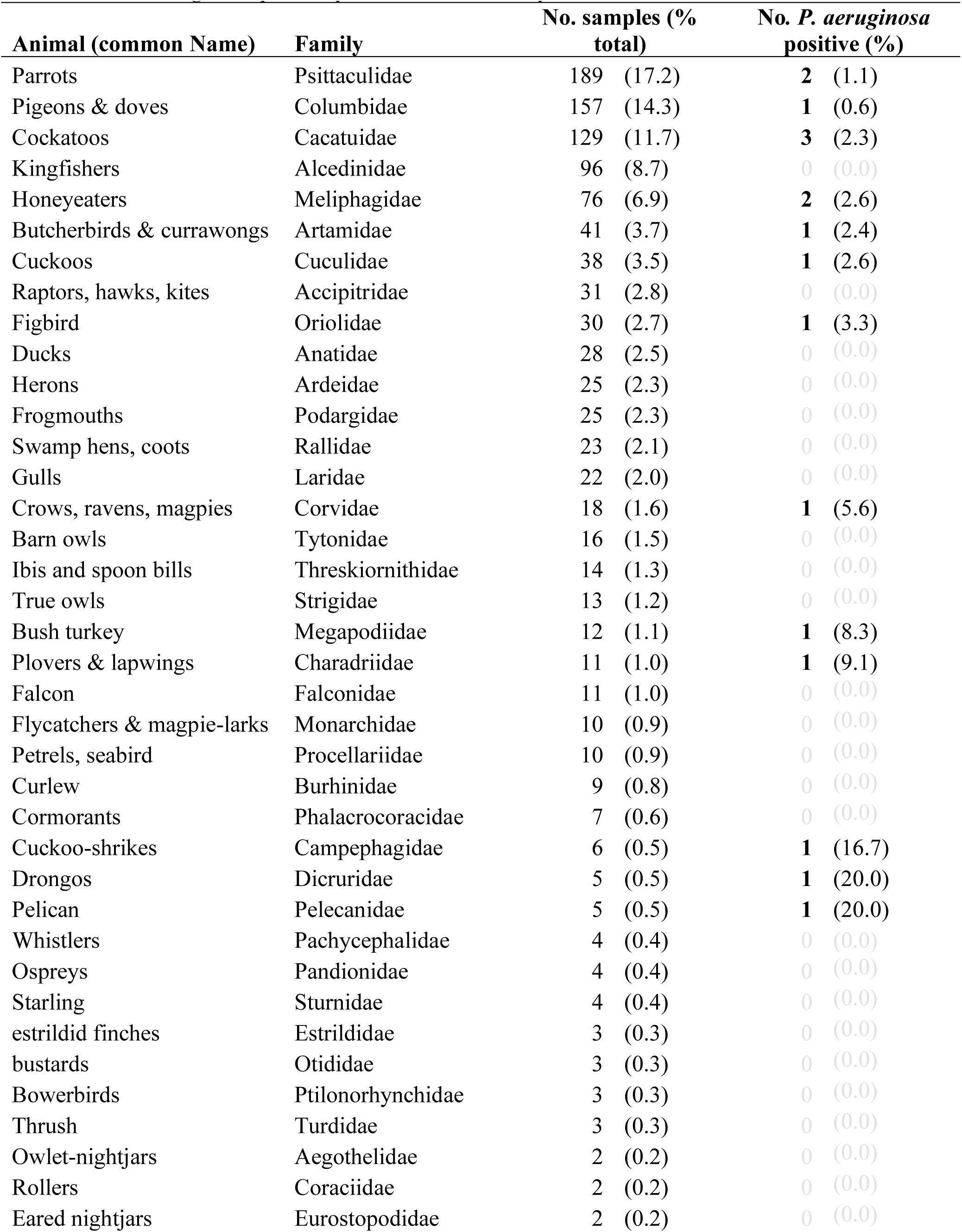
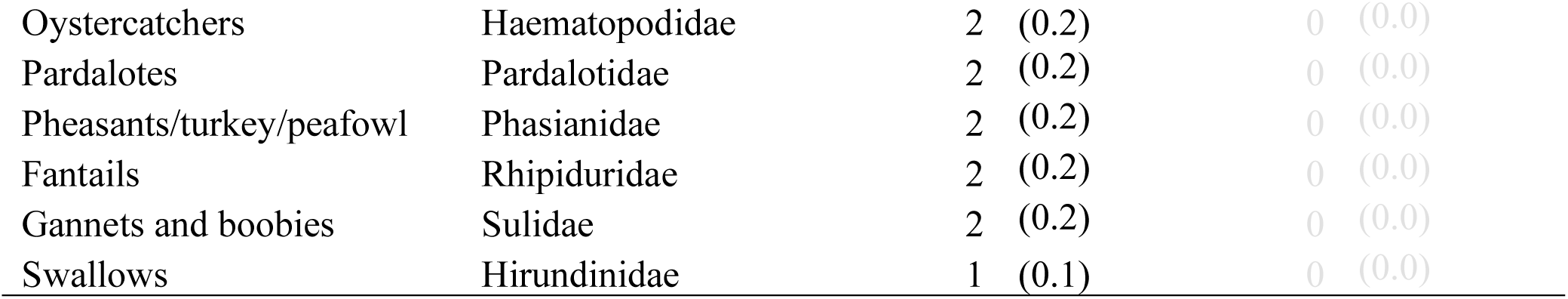
Bird Families identified from 1,101 wild bird DNA samples and corresponding*Pseudomonas aeruginosa* positivity rates for each Family.

**Table S2.** Anatomical site swabbed for each retrospective DNA sample obtained and corresponding Pseudomonas aeruginosa positivity rate.

| **Animal** | **Sample type** | **No. samples** | ***P. aeruginosa* positive (%)** |  |
| --- | --- | --- | --- | --- |
| Wild Birds (*n*=565) | Total | 1101 | 17 | (1.5) |
|  | Eye | 565 | 16 | (2.8) |
|  | Cloaca | 534 | 1 | (0.2) |
|  | Other | 2 | 0 | (0.0) |
| **Domestic Animals** |  |  |  |  |
| Cats (*n*=138) | Total | 139 | 3 | (2.2) |
|  | Eye | 5 | 1 | (20.0) |
|  | Faeces | 130 | 1 | (0.8) |
|  | Nose | 2 | 0 | (0.0) |
|  | Mouth | 1 | 1 | (100.0) |
|  | Rectum | 1 | 0 | (0.0) |
| Dogs (*n*=126) | Total | 126 | 1 | (0.8) |
|  | Eye | 2 | 0 | (0.0) |
|  | Faeces | 112 | 1 | (0.9) |
|  | Fur | 4 | 0 | (0.0) |
|  | Joint fluid | 4 | 0 | (0.0) |
|  | Mouth | 1 | 0 | (0.0) |
|  | Prepuce | 1 | 0 | (0.0) |
|  | Uterus | 1 | 0 | (0.0) |
|  | Vagina | 1 | 0 | (0.0) |
| Guinea pig (*n*=1) | Total | 3 | 0 | (0.0) |
|  | Eye | 1 | 0 | (0.0) |
|  | Mouth | 1 | 0 | (0.0) |
|  | Vagina | 1 | 0 | (0.0) |
| Pet Bird (*n*=1) | Total | 1 | 1 | (100.0) |
|  | Eye | 1 | 1 | (100.0) |
| Rabbit (*n*=1) | Total | 1 | 0 | (0.0) |
|  | Eye | 1 | 0 | (0.0) |
| **Livestock** |  |  |  |  |
| Horses (*n*=68) | Total | 68 | 5 | (7.4) |
|  | Faeces | 33 | 1 | 3.0 |
|  | Cervix | 2 | 0 | (0.0) |
|  | Clitoris | 3 | 0 | (0.0) |
|  | Penis | 2 | 2 | (100.0) |
|  | Nose | 8 | 0 | (0.0) |
|  | Nasal-vaginal pooled | 13 | 0 | (0.0) |
|  | Respiratory swab | 2 | 0 | (0.0) |
|  | Tracheal wash | 2 | 2 | (100.0) |
|  | Uterus | 2 | 0 | (0.0) |
|  | Vagina | 1 | 0 | (0.0) |
| Cattle (*n*=45) | Total | 45 | 0 | (0.0) |
|  | Vagina | 34 | 0 | (0.0) |
|  | Rectum | 10 | 0 | (0.0) |
|  | Joint fluid | 1 | 0 | (0.0) |
| Goat (*n*=1) | Total | 14 | 0 | (0.0) |
|  | Brain | 1 | 0 | (0.0) |
|  | Eye | 1 | 0 | (0.0) |
|  | Intestine | 1 | 0 | (0.0) |
|  | Joint fluid | 2 | 0 | (0.0) |
|  | Liver | 1 | 0 | (0.0) |
|  | Lung tissue | 6 | 0 | (0.0) |
|  | Rectum | 1 | 0 | (0.0) |
|  | Spleen | 1 | 0 | (0.0) |
| Alpaca (*n*=1) | Total | 3 | 1 | (33.3) |
|  | Eye | 1 | 0 | (0.0) |
|  | Rectum | 1 | 1 | (100.0) |
|  | Vagina | 1 | 0 | (0.0) |
| Poultry (*n*=1) | Total | 3 | 0 | (0.0) |
|  | Mouth | 1 | 0 | (0.0) |
|  | Cloaca | 1 | 0 | (0.0) |
|  | Eye | 1 | 0 | (0.0) |
| **Native wild marsupials** |  |  |  |  |
| Kangaroos (*n*=39) | Total | 39 | 0 | (0.0) |
|  | Faeces | 39 | 0 | (0.0) |
| Koala (*n*=67) | Total | 127 | 2 | (1.6) |
|  | Eye | 48 | 1 | (2.1) |
|  | Nasal | 22 | 1 | (4.5) |
|  | Urogenital tract | 45 | 0 | (0.0) |
|  | Oropharynx | 3 | 0 | (0.0) |
|  | Rectum | 7 | 0 | (0.0) |
|  | Endotracheal tube | 1 | 0 | (0.0) |
|  | Pleural cavity | 1 | 0 | (0.0) |

*n* = number of individual animals.

